# Unconventional *IFNω*-like genes dominate the type I IFN locus and the constitutive antiviral responses in bats

**DOI:** 10.1101/2024.02.22.581518

**Authors:** Rong Geng, Qi Wang, Yu-Lin Yao, Xu-Rui Shen, Jing-Kun Jia, Xi Wang, Yan Zhu, Qian Li, Zheng-Li Shi, Peng Zhou

**Author notes:** Correspondence (P.Z.) or (Z.L.S.).

## Abstract

Bats are the natural reservoir hosts of some viruses, some of which may spillover to humans and cause global-scale pandemics. Different to humans, bats may coexist with high pathogenic viruses without showing symptoms of diseases. As one of the most important first defenses, bat type I interferon (IFN-Is) were thought to play a role during this virus coexistence and thus were studied in recent years. However, there are arguments that whether bats have a contracted genome locus or constitutive expressed IFNs, mainly due to species-specific findings. We hypothesized that because of the lacking of pan-bat analysis, the common characters for bat IFN-Is have not been revealed yet. Here, we characterized the IFN-I locus for 9 Yangochiroptera bats and 3 Yinpterochiroptera based on the their high quality bat genomes. We also compared the basal expression for 6 bats and compared the antiviral, anti-proliferative activity and thermo-stability of a representative *Rhinolophus* bat IFNs. We found a dominance of unconventional *IFNω*-like responses in the IFN-I system, which is unique to bats. In contrast to *IFNa*-dominated IFN-I loci in the majority of other mammals, bats generally have shorter IFN-I loci with more unconventional *IFNω*-like genes (*IFNω* or related *IFNaω*), but with less or even no *IFNa* genes. In addition, bats generally have constitutively expressed IFNs, the highest expressed of which is more likely an *IFNω*-like gene. Likewise, the highly expressed IFNω-like protein also demonstrated the best antiviral activity, anti-proliferative activity or thermo-stability, as shown in a representative *Rhinolophus* bat species. Overall, we revealed pan-bat unique characteristics in IFN-I system, which provide insights into our understanding of the innate immunity that contribute to a special coexistence between bats and viruses.

## Introduction

Bats are the natural reservoir hosts of viruses. Some of the bat-origin viruses, such as Ebola virus or SARS-related coronaviruses may be highly pathogenic to humans or causing global-scale pandemics (1). However, as the natural reservoir hosts, bats may coexist with these viruses without showing symptoms of diseases (2). During the past several years, both our research and that of others have revealed distinctive bat characteristics that may contribute to this bat-virus coexistence (3–6). Yet, we are still far from a full understanding of bat immunity.

It is believed that the unique flight ability of bats, as the only flying mammals, has significantly influenced the shaping of their genes, particularly those associated with metabolic and innate immune pathways (6). As the first line of antiviral defense, type I interferons (IFN-I) play a vital role in controlling viral replication. In non-bat mammals, multiple copies of *IFNa*-like genes dominate the type I IFN locus, followed by species-specific IFN genes (*IFN*-*ω*, -*τ*, *δ* and other), whereas single copied *IFNβ* and *IFNε* normally determined the boundary of type I IFN locus in genome. Previously, we characterized bat type I IFN system and demonstrated a contracted IFN-I locus but constitutively expressed *IFNa* and related IFN-stimulated genes (ISGs) in *Pteropus alecto* bat, showed a “less is more” characteristic of bat IFN system (3). However, a later study demonstrated an expanded IFN-I locus in *Rousettus aegyptiacus* without a basal IFN expression (7). These species-specific features confounded our understanding of the true nature of pan-bat IFN-I responses (NIH R21). On one hand, both *Pteropus* and *Rousettus* bats are Yinpterochiroptera (mostly frugivores), and the IFN characteristics of Yangochiroptera (mostly insectivorous) are still unknown (8). On the other hand, we believe the fundamental properties of bat IFN-I system have not been revealed yet, largely due to the absence of a comprehensive pan-bat analysis.

Here, we hypothesize there are pan-bat characteristics within the IFN-I systems that are distinct from other mammals. Based on the recent high quality bat genomes, we characterized IFN-I locus in 12 bats, including 9 Yangochiroptera bats and 3 Yinpterochiroptera. Additionally, we conducted a comprehensive analysis, comparing the basal expression profiles, the antiviral functionality and thermo-stability for representative bat IFNs. We found a dominance of unconventional *IFNω*-like responses in bat IFN-I systems, which is very different from all other mammals. The study may shed light on our understanding on a special coexistence between bats and viruses.

## Materials and Methods

### Isolation of primary cell and cell culture

Healthy *Rhinolophus ferrumequinum* bats from Jiyuan City, Henan Province, were captured and anaesthetized with isoflurane. The animal work was approved by the Ethics Committee of the Wuhan Institute of Virology (WIVA43202105). Various organs were isolated, washed with HBSS (14025092, Gibco), and cut into small pieces. After washing with 5 mL HBSS, the organ pieces underwent digestion with 10 mL 0.25% trypsin-EDTA (25200-072, Gibco) at 4 °C overnight. Following a 24-hour digestion period, cells were further digested at 37 °C for 1 hour with oscillated every 10 minutes. The digested cells were filtered through a 70µm strainer, centrifuged at 1000 rpm for 5 minutes. The primary cells were resuspended in RPMI 1640 medium (A10491-01, Gibco) with 20% fetal bovine serum (FBS; Gibco, 10099-141C) and cultured at 37 °C with 5% CO_2_ .

HEK293T-sg cells were cultured with DMEM + 10% FBS (Gibco) at 37 °C in 5% CO_2_ for the package of lentivirus encoding SV40LT (plasmids provided by Dr. Lin-Fa Wang, Duke-NUS Medical School). To establish immortalized cell lines, primary *Rhinolophus ferrumequinum* bat cells were seeded in 6-well plate at a density of 6×10^5^ cells/well with DMEM containing 10% FBS (Gibco). After 12 h, the culture medium was replaced with the lentivirus described above. 48h later, the culture medium was replaced with fresh medium with 10 µg/ml of hygromycin (10687010, Thermo Scientific) for the selection of SV40LT-positive cells. The RfKT cells, originating from immortalized *Rhinolophus ferrumequinum* bat kidney origin cells were utilized in the study, along with RfBT (brain), RfPT (pulmonary) or RfHT (heart) cells.

### Interferon genome locus Analyses

The identification and annotation of interferon gene loci were conducted in three sequential steps. Initially, *Pteropus alecto* IFN-I loci (9) were designated as reference genes, and various bat genomes were searched for their respective type I interferons utilizing blastn v2.6.0. Sequence between *IFNε* and *IFNβ* was considered as candidate loci. To avoid missing genes, 1M bp upstream and downstream of the candidate locus were also analyzed. Subsequently, all potential open reading frames (ORFs) with the locus were identified using ORFfinder v0.4.3, and interferon-related ORFs were selected using HMMER hmmscan v3.1b2 against the Pfam database. Finally, functional interferons with the intact domains predicted by SMART were selected. Following annotation, all bat IFN-Is were employed as reference genes, and blastn was conducted again to check for any potentially missed IFNs.

Sequences alignments of the IFN-Is gene in this study were performed using MEGA11 and BioEdit (v7.1.3.0). A phylogenetic tree was constructed utilizing a maximum likelihood (ML) method with the HKY substitution model by IQ-TREE (v1.6.1), and visualized using FigTree v1.4.4 (tree.bio.ed.ac.uk/software/figtree/). Bootstrap confidence values determined through 1000 replicates.

### RNA-seq data analysis

RNA-seq data in fastq format was aligned to the reference genome utilizing HISAT2 v2.2.1 (10). The resulting intermediate SAM files were sorted by SAMtools v1.6 and subsequently input into StringTie v2.1.7 (11) for transcriptome assembly and estimation. A new annotation of interferons was added to GTF file for transcriptome estimation. Package DESeq2 v1.36.0 (12) in R v4.2.1 was employed for normalizing expression matrix and comparing the expression level of genes. Log2 fold change greater than 2 and p-value less than 0.05 was identified as differential expression gene.

### Validation of Rf-IFNα/w binding to IFN-αR

Knockdown (KD) of the *Rf-IFNAR1* gene was achieved by transducing of RfPT cells with lentiviruses expressing specific siRNA (Rf-IFNaR1: 5’- GCCTGACTGTCAATATATTAC - 3’). Subsequently, the transduced cells were cultured with puromycin (8 µg/ml) for 7 days. Due to the the unavailability of a suitable antibody for *Rhinolophus ferrumequinum* IFNaR1, the validation of the KD by western blot was not feasible. Consequently, we examined the expression of *IFNAR1* gene by qPCR to confirm IFNaR1-KD. Both wild-type RfPT and IFNaR1 knockdown cell lines were treated with 1000 ng/ml *Rf*-IFN proteins for 24 h. The cells were then harvested and lysed, and subjected to western blotting using the anti-Human phospho-STAT1 (1:1000, Y701, CST). Beta-tubulin was used as an internal control in the WB (1:5000, 66240-1-Ig, Proteintech).

Ruxolitinib (Selleck, S1378) was diluted to a final concentration of 100 mM, and 200 µl (for a 24-well plate) of Ruxolitinib solution was added to cells. One hour later, cells were washed twice with PBS and treated with different dilutions of *Rf*-IFN proteins. After 24 h, the expression of cellular p-STAT1 protein was determined by western blot.

### Rhinolophus ferrumequinum bat IFN protein expression

The ORFs of *Rf* bat IFN were inserted into pCAGGS vectors with an N-terminal S-tag. The sequences of the three types of IFNs were as follows: *IFNa* (XM_033123262.1), *IFNω* (XM_033124139.1), and *IFNaω* (XM_033123266.1). Constructed plasmids were transiently transfected into HEK293F (mammalian expression system). The supernatant collected for protein purification was purified with Centricon® centrifugal filter (10 K NMWL, Millipore). Concentrated IFN proteins were verified by western blot.

### Antiviral Activity Test

Herpes Simplex Virus type 1 (HSV1-GFP, generously provided by Yanyi Wang and Minhua Luo, Wuhan Institute of Virology, Chinese Academy of Sciences), and PR8 (H1N1 influenza A virus strain, a kind gift from Quanjiao Chen, Wuhan Institute of Virology, Chinese Academy of Sciences) were used in this study. Briefly, cells were seeded to 48-well plate and grown to more than 90% confluence. *Rf* bat cells were incubated with 10^-3^ µg/mL of *Rf*-IFN proteins at 37 °C for 6, 9, and 12 hours. Subsequently, the cells were infected with HSV1 or PR8 (MOI=0.5) and samples were collected at 24 hours post-infection (hpi). Then *Rf* bat cells were treated with 1:10 serially diluted *Rf*-IFN proteins at concentrations ranging from10^-3^-10^-7^ µg/ml at 37 °C before infection. Six hours later, cells were infected with HSV1 or PR8 at the indicated MOI and samples were harvested at 24 hpi. Viral load was determined by qRT-PCR or immunofluorescence assay. The gene specific primers for qPCR were shown in Table S1.

### Thermo-stability Test

To examine the effect of temperature on the *Rf*-IFN proteins, RfKT cells were seeded in 24-well plate at a density of 4 × 10^5^ cells/well. *Rf*-IFN proteins at a concentration of 1 × 10^-3^ µg/ml were incubated at 4 °C or 42 °C for 4 hours. Subsequently, the proteins were added to the cells and cultured at 37 °C. After 6 hours, the cells were infected with HSV1 and PR8 (MOI = 0.5), and samples were collected at 24 hpi. Viral RNA in the cytoplasm was determined by qPCR.

### Anti-proliferative activity assay

Cell Counting Kit-8 (HY-K0301-3000T, MCE) was used for assessing cellular viability. Twenty thousand cells in 100 µl volume were added to a 96-well plate in triplicate. RfKT cells were treated with or without IFN proteins (1 ng/ml for each) at 37 °C for 0, 24, 48, 72 h. 10 µl of CCK-8 reagent were added to every well and incubated for 2 h at 37 ℃. Absorption was measured at 450 nm using an M200 Pro (TECAN) spectrometer to assess the activity of the cells. Cell activity = [(As-Ab) / (Ac-Ab)] × 100%, where As is the experimental hole absorbance , Ac is the control hole absorbance and Ab is the blank hole absorbance.

BrdU (5-bromo-2’-deoxyuridine) incorporation assay was also used to evaluate the proliferative rate of cells. Briefly, cells were seeded to 24-well plate with three million cells per well. Following the treatment with IFN proteins for 48 hours as described above, cells were exposed to 10 µM of BrdU. After 1 later, cells were washed three times with PBS and fixed with cold 70% methanol for 1 h, followed by blocking with 5% BSA at room temperature for 30 min. Then 1.5M of HCl was added to cells and incubated at room temperature for 30min. BrdU that inserted to cell DNA was measured by anti-BrdU antibody (1:500, 66241-1-Ig, Proteintech) and a mouse anti-Cy3 antibody (1:100, ab6939, Abcam) to evaluate the cell proliferation rate. Cell nuclei were stained with DAPI (Beyotime, C1002). Staining patterns were examined using Fluorescence imaging system on an Invitrogen™ EVOS™ M5000 microscope (ThermoFisher).

### RNA extraction and qRT-PCR

RNA from both host and virus was extracted by using the RNAsimple Total RNA Kit (DP419, TIANGEN) and Virus DNA/RNA Extraction Kit (RM401-04, Vazyme) respectively according to the manufacturer’s instruction without modification. Host nucleic acid were used for RNAseq analysis. RT-qPCR was conducted using HiScript II One Step qRT-PCR SYBR Green Kit (Q221-01, Vazyme) with gene-specific primer pairs on a CFX Connect Real-Time PCR Detection System (Bio-Rad Laboratories). The thermal cycling protocol was as follow: one cycle at 50 °C for 15 min, 95 °C for 30 s, 40 cycles of 95 °C for 10 s, and 60 °C for 30 s. The beta-actin gene was used for the normalization of the housekeeping gene. Primers used for qPCR detection in this study were listed in Table S1.

### Immunofluorescence assay

Untreated or IFN-treated cells of RfKT and RfBT were infected with HSV1-GFP or PR8 (MOI=1) for 24 h. Cells were then fixed and permeabilized. For PR8 infected cells, mouse-anti-NP (1:1000, sc-101352, Santacruz) was applied for 1 hour at room temperature. After washing three times with PBS, cells were incubated with mouse anti-Cy3 (1:500, ab6939, Abcam) at room temperature for 2 h. Cell nuclei were stained with 5 µg/ml of DAPI (10236276001, Roche Diagnostics).

### Statistical analysis

Data analyses were performed using GraphPad Prism 8.3 software. Data were shown as mean ± standard error of mean (SEM). Statistical analysis was determined using one-way ANOVA Dunnett’s T3 test. P values < 0.05 were considered statistically significant.

## Results

### Identification of a pan-bat contracted IFN-I locus and a Yin-/Yang-bat difference

To fully understand the IFN-I systems in bats, we initiated our investigation by analyzing the genomic locus. Recognizing the impact of genome quality on locus annotation, we only selected high-quality genomes (9), including 9 Yangochiroptera bats, 3 Yinpterochiroptera bats, as well as 8 non-bat mammals representing diverse evolutionary stages. We re-analyzed *Pteropus alecto* and *Rousettus aegyptiacus*, two bat species previously reported (3, 7), based on their newly updated genomes. Our analysis further encompassed a reservoir host of SARS-related coronavirus (*Rhinolophus ferrumequinum*), and the species that may serve as intermediate host of bat coronaviruses, pangolin (*Manis javanica*) (13, 14). The putative functional or pseudotyped IFNs were shown (Figure 1 and Figure S1).

**Figure 1.**
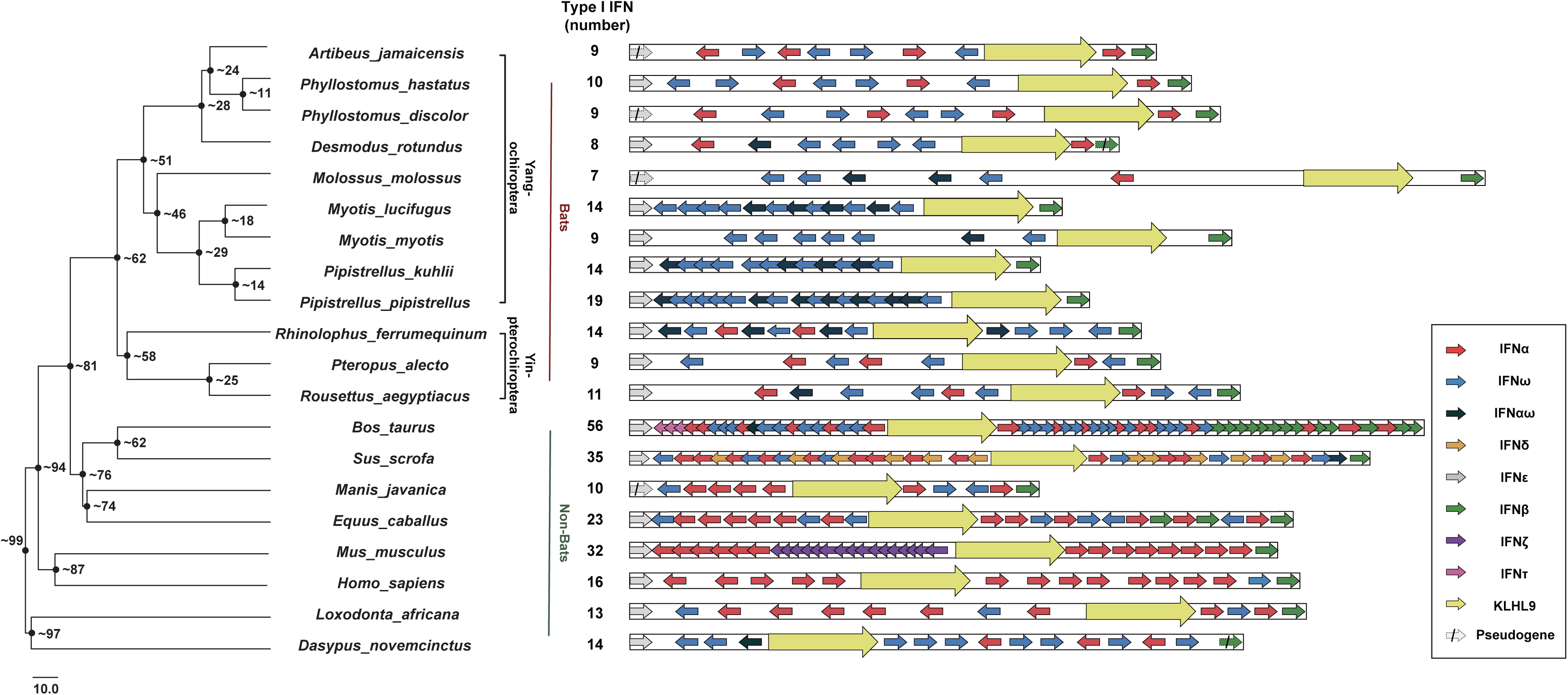
The type I IFNs (IFN-Is) genome locus in bats and non-bat mammals. Genome locus and annotation for 12 bats (9 Yang- and 3 Yin-) and 8 non-bat mammals representing different evolutionary stages. The genome information can be found in Table S2. The length of the locus was drawn to scale. The solid arrows represent IFN ORFs and dotted arrows represent pseudogenes, all with directions. The number of IFN-Is (excluding *KLHL9*) genes for each species is shown on the right. The phylogenetic tree is drawn according to timetree (timetree.org), and the estimated divergence times are denoted (time, million years).

Our first observation is a general shorter IFN-I genome locus with lower IFN-I gene numbers in bats under analysis (Figure 1 and Table S2). Most bats have 7-14 IFN genes, and the only exception is *Pipistrellus pipistrellus* bat (19 genes). This number is lower compared to many evolutionarily related mammals, such as cow and pig. Notably, the analysis resulted in only 11 IFN-I genes in *Rousettus aegyptiacus* bat, a number that is much lower than previously reported (7). It is hypothesized that the genomic size of the IFN locus is expanding following evolution as gene duplication has occurred in the vertebrate IFN-I in a step-wise manner (15). However, bats and pangolins are the two exceptions of the species analyzed here in general. Furthermore, we also observed dramatic differences between Yangochiroptera and Yinpterochiroptera bats. The IFN-I locus was normally separated into two sections by a non-IFN gene *KLHL9*, with the IFN genes in each section expanding independently (15). As all other mammals and the 3 Yinpterochiroptera bats keep both sections, the 9 Yangochiroptera bat species almost lost the section between *KLHL9* and *IFNβ* entirely (Figure 1). To find out whether the reduced IFN-I gene number results from pseudogenization during gene expansion, we compared the pseudogenes in IFN-I locus between bats and other species. The data indicated fewer IFN pseudogenes in the bats species under analysis, suggesting a lack of IFN-I gene expansion at the first beginning (Figure S1). This implies that bats appear to choose a different IFN-I evolutionary path to keep only limited numbers of genes, as hypothesized previously (3). Additional supporting evidence includes the pseudogenization of *IFNβ* or *IFNε* genes, with at least one of the two single copied IFN genes were lost in four bats. Similar losses were only observed in pangolin (*IFNε*) and armadillo *(IFNβ*) (Figure 1 and Figure S2).

### General dominance of unconventional IFNω-like genes in bat type I IFN locus

Our next finding is the general dominance of *IFNω*-like genes in bat type I IFN locus (Figure 1 and 2). In the evolutionary analysis of vertebrates IFN-I locus, it is believed that *IFNβ* and *IFNε* were the first genes to evolve, followed with a subsequent *IFNω*- like duplicate called *IFN-αω*. From here, this *IFN-αω* may evolve into *IFN-α* types or *IFN-ω* types in a random profile, whereas the rather ancient *IFN-αω* gene type was lost in majority of mammals (15). Unlike the majority of placental mammals choose *IFNα* over *IFNω* genes, bats choose a different evolutionary path. The first evidence is that bats generally have more *IFNω* genes than *IFNα* genes. An extreme example is the entire lost of *IFNα* gene in the *Myotis* and *Pipistrellus* bat species, a pattern that is not found in other mammals. The next evidence is the general reservation of the ancient *IFNαω* gene. This gene can be found in 8 out of 12 bat species studied, and the gene number ranges from 1-7 copies. As comparison, only one copy of *IFN-αω* gene can be identified in armadillo, pig or cow, but not in other mammals analyzed (Figure 2A) (15). From ORFs or pseudogenes analysis, it is suggested that *IFNαω* gene tends to resemble *IFNω* more than *IFNα* genes (15). Thus, we conclude that *IFNω*-like genes dominate bat IFN-I loci, which pattern is similar to armadillo, a rather ancient placental mammal. Thus, bat IFN-I loci appear to be evolutionarily ancient (Figure 2B and 2C).

**Figure 2.**
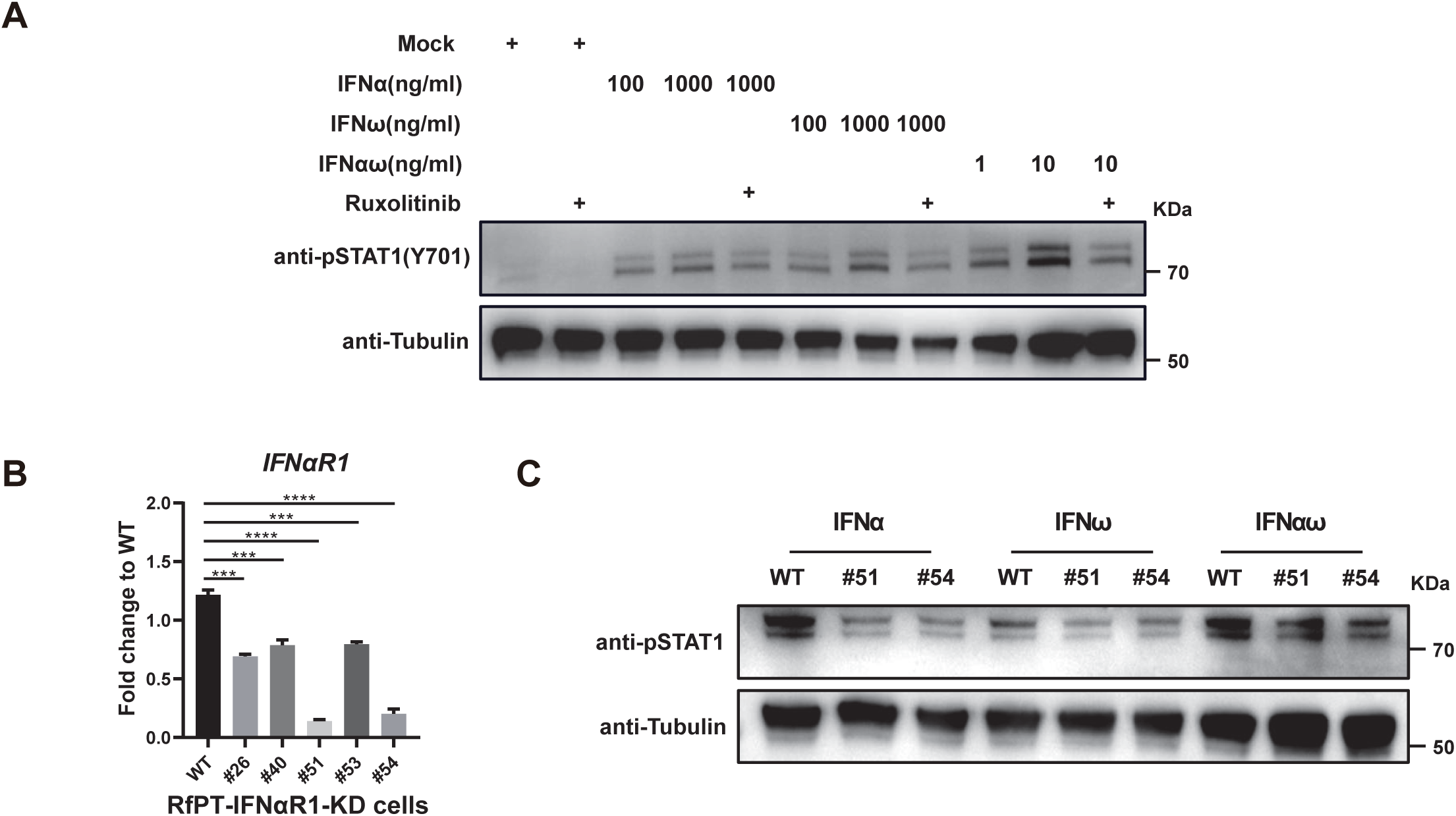
The evolution of bat IFN-I genes. (**A**) Statistics for each IFN-I subtypes in bats and non-bat mammals, specifically on *IFNa*, *IFNω* and *IFNaω* genes. (**B**) Phylogenetic analysis of bat IFN-I genes. All IFN-I subtypes from 12 bat species (colored) and armadillo (black) were subjected for phylogenic analysis using maximum likelihood method, with IFN-III gene from human (red) as outgroup. (**C**) The proposed evolutionary path for IFN-I genes in bats and non-bat mammals. (D) The maps show the global geographic distribution of bats in this study based on publicly available data from the IUCN Red List of Threatened Species Database as of January 20, 2023 (iucnredlist.org).

### IFNω-like genes dominate the basal expression in bats

In a previous study, we showed the constitutive mRNA expression of *IFNα* genes in *Pteropus alecto* bat, which pattern was then confirmed at the protein level and in other bats (3, 16, 18). Here, we first performed a gene expression analysis of all IFN-I gene types to a list of bat species using published tissue expression data (*Pteropus*, *Rousettus*, *Myotis*, *Pipistrellus* and *Molossus* bats) (9) or experimental data by ourselves (*Rhinolophus* bats) (Table S3). The geographical distribution of these bats spans major continents, representing the major bat population in the world (Figure 2D). Due to the high sequence similarity, we only detected the sum up expression of each IFN gene types, specifically *IFNε*, *IFNβ, IFNaω*, *IFNω* or *IFNa*. The data indicated a general basal expression of at least one IFN-I gene in bats, with the specific type being *IFNaω*, *IFNω* or *IFNa* depends on the species or tissue. In most tissues, *IFNω*-like genes showed the highest basal expression. This finding is consistent with our previous observations, where *IFNa* was constitutively expressed in *Pteropus alecto* but at a relatively lower level than *IFNω* (Figure 3A). There are 2 *IFNa*, 4 *IFNaω* and 6 *IFNω* gene copies in the IFN-I locus for *Rhinolophus ferrumequinum* bat (Figure S3). The qPCR data indicated a basal mRNA expression of *IFNω*-like genes in all tissues detected, whereas *IFNaω* is the highest expressed IFN in the two immortalized *Rhinolophus* kidney (RfKT) or brain (RfBT) cell lines (Figure 3B). However, *IFNaω* is also the least induced IFN following HSV1 and Influenza virus infection (Figure 3C). Overall, we found a pan-bats constitutive expression of *IFNω*-like genes, suggesting a potentially more significant role for this gene type in bats compared to most other mammals.

**Figure 3.**
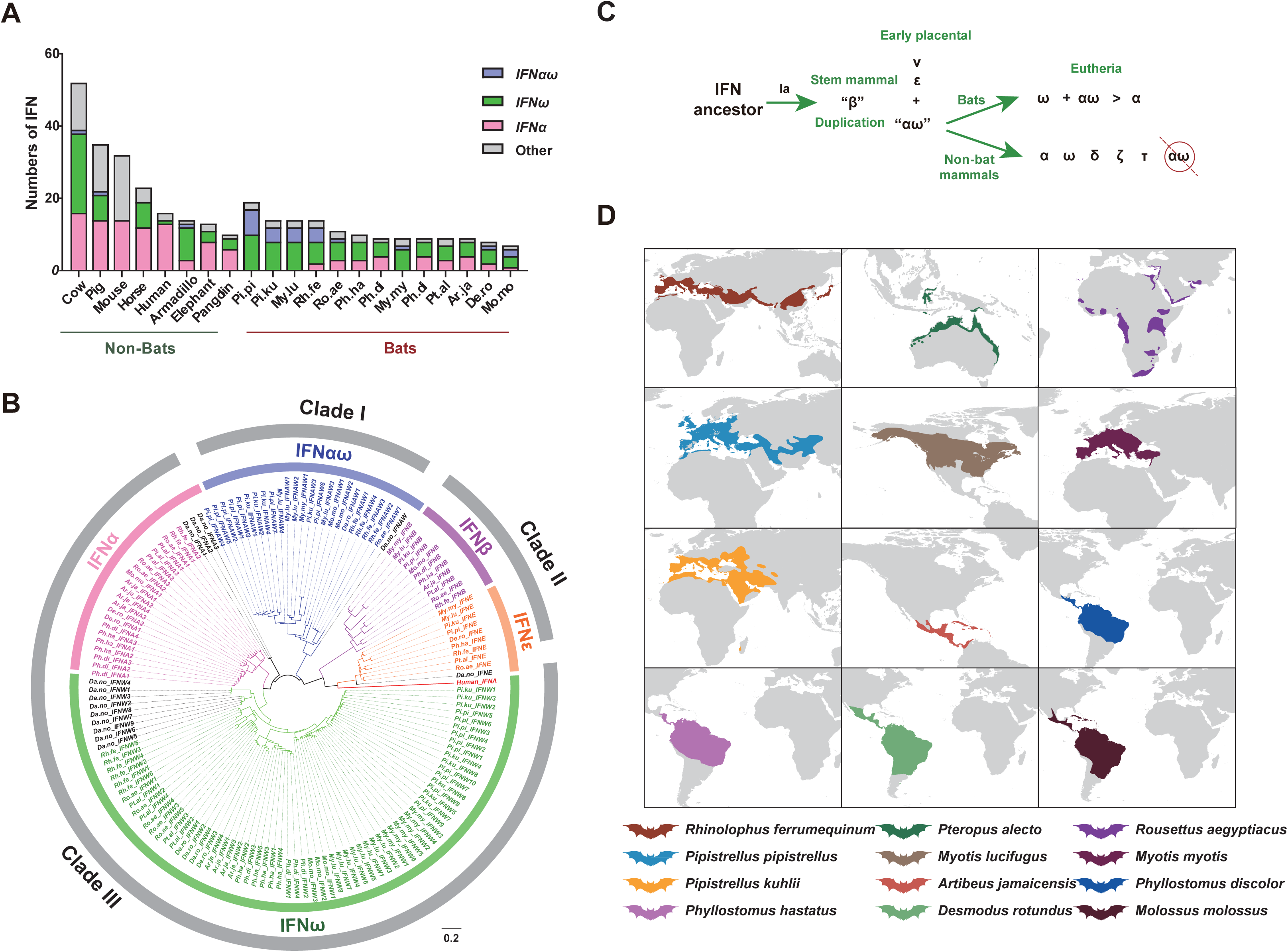
Constitutive IFN-I gene expression for bats. (**A**) Boxplots shows the baseline expression of IFN-I subtypes in 5 bat species. Analysis is based on public tissue RNA-seq data (Table S3). (**B**) The baseline expression of IFN-I subtypes in tissues and cells (kidney cells RfKT) from *Rhinolophus ferrumequinum* bat. Relative gene expression was analyzed using qPCR. The Ct values were normalized with a housekeeping gene (beta-actin). (**C**) IFN-I genes upregulated during viral infection. RfKT cells were infected with HSV1 or Influenza virus PR8 at an MOI of 0.5, and samples were collected at 24 h post infection. Gene expression was determined by qPCR. Data are means ± SEM; n = 3 replicates of 2–3 independent assays, the same letter indicates no significant difference, while the different letter indicates significant difference (P = 0.05). Mean values were analyzed using one-way ANOVA with Tukey’s correction. Compared each column to every other column.

### IFNαR1 is the functional receptor for IFN-Is in Rhinolophus bat

To test whether IFN-Is in *Rhinolophus ferrumequinum* bats still use IFNaR1 as functional receptor, we employed Ruxolitinib, a potent and selective JAK1/2 inhibitor targeting the JAK/STAT signaling pathway activated by both type I and type III interferon. Cells were pre-treated with Ruxolitinib for 1 h before stimulate with Rf-IFN proteins, which resulted in a significant inhibition of p-STAT1 downstream of the interferon pathway, as evidenced by western blot results (Figure 4A). This inhibition indicated the regulation of these three IFN-Is by the JAK/STAT pathway. Similarly, we constructed RfPT-IFNaR1 knockdown cell clones. Intracellular expression of IFNaR1 RNA was determined by qPCR, and clones #51 and #54 were selected for subsequent experiment (Figure 4B). Following incubation with Rf-IFN proteins for 24 hours, the results demonstrated a decrease in p-STAT1 in knockdown cells compared to WT cells (Figure 4C). These findings strongly suggest that these IFN-Is still use IFNaR1 receptor in *Rhinolophus ferrumequinum*.

**Figure 4.**
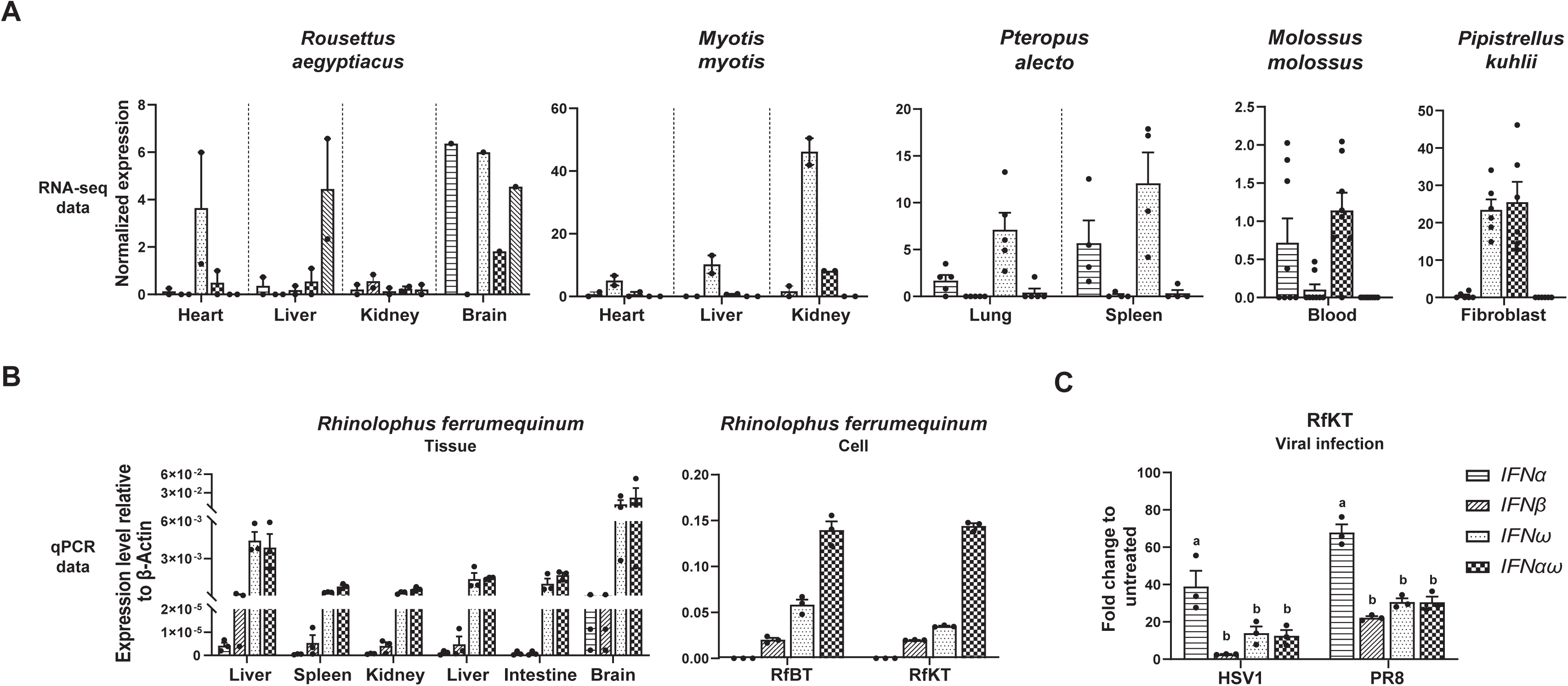
Rf-IFNa/ω signaling rely on the IFN-aR. (**A**) Effect of IFNs and Ruxolitinib on RfPT cells. RfPT cells were incubated with the indicated doses of Rf-IFN proteins and Ruxolitinib (added 1 h before IFNs). 24 h later, anti-pSTAT1 (1:1000, Y701, CST) and beta-tubulin (1:5000, 66240-1-Ig, Proteintech) were analyzed using immunoblotting. Molecular markers were shown on the right sides of the blots. (B) The expression level of IFNaR1 in RfPT-IFNAR1-KD cells, different monoclonal cells were analyzed using qPCR. (C) The effect of IFNaR1 knockdown on interferon response. RfPT-IFNaR1-KD cells were treated with the indicated Rf-IFN proteins, and the indicated antibodies were detected by western blot after 24 h. The data are representative of three independent experiments. Statistical analyses were carried out using one-way ANOVA.

### IFNαw protein showed the most potent antiviral functionality in Rhinolophus bat in vitro

To understand the biological relevance of IFN-I gene types and their mRNA expression levels, we conducted a comparative analysis of the functionality of IFN-I genes, focusing on *Rhinolophus ferrumequinum* bat. A representative gene for *IFNaω*, *IFNω* or *IFNa* were subjected for protein expression, followed by an evaluation of antiviral, ISG induction, thermo-stability and anti-cell proliferation analysis.

In the IFN antiviral functional test, the RfKT cells were treated with serially diluted purified *Rf* bat IFN proteins, followed by infection with HSV1 or Influenza virus PR8. The viral replication level was determined by qPCR targeting at viral RNA. All three IFNs demonstrated antiviral activity in a dose-dependent and time-dependent manner, whereas IFNaω showed the highest activity. The IFNa protein displayed the weakest against the two viruses (Figure 5A). The same conclusion can be determined using additional three *Rf* bat cell lines (Figure 5B). Viral replication levels were also determined using IFA by measuring viral infection induced syncytia or percentage of virus infected cells (Figure 5C and 5D). Similarly, IFNaω demonstrated that highest antiviral activity without cell differences.

**Figure 5.**
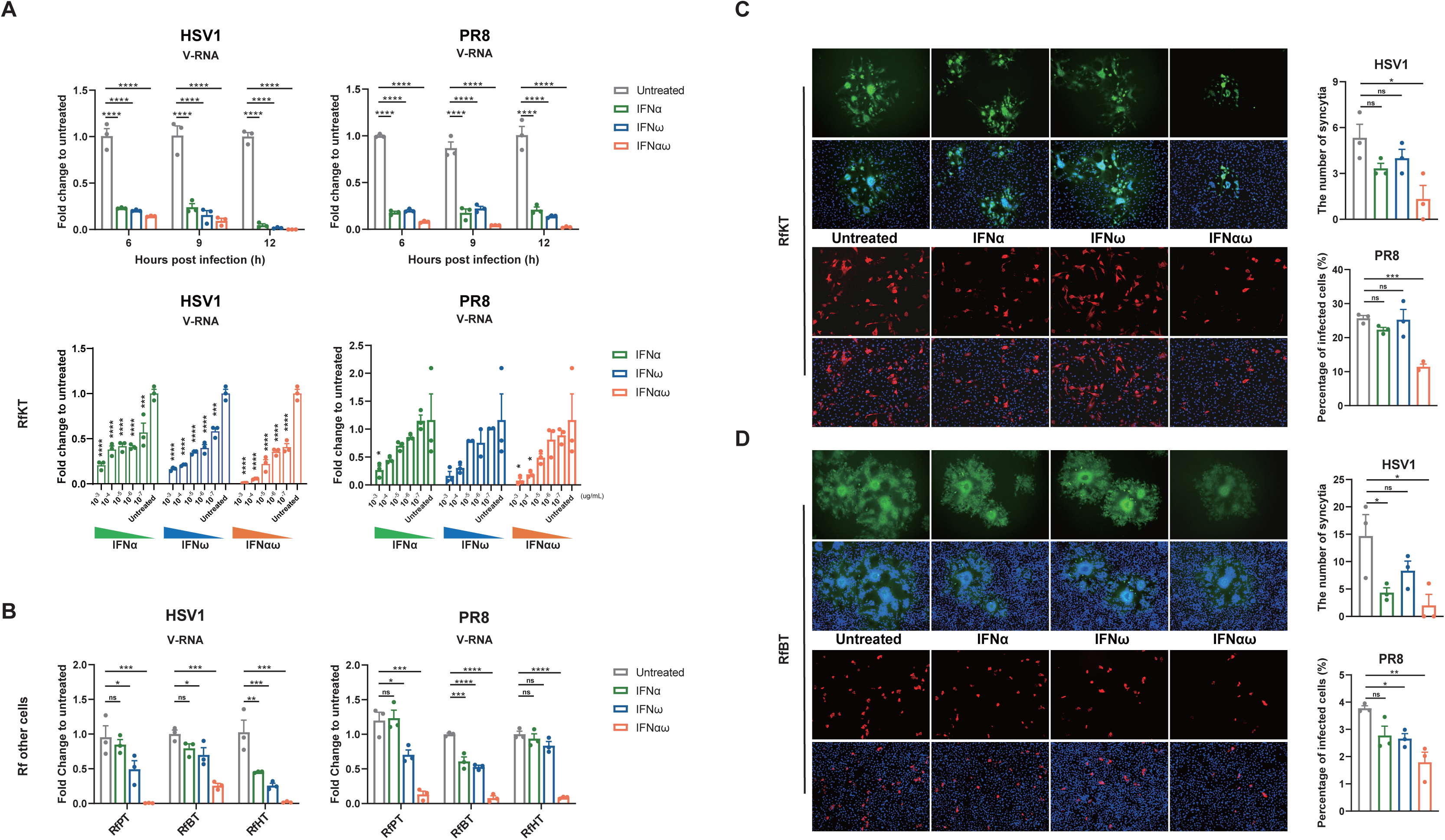
The antiviral activities for *Rhinolophus ferrumequinum* bat IFN-Is. (**A**) RfKT cells were incubated with Rf-IFN proteins (10^-3^ µg/ml) at 37 ℃ for 6, 9, 12 hours, then infected with HSV1 or PR8 (MOI=0.5). RfKT cells were treated with different dilutions of *Rhinolophus ferrumequinum* IFN proteins (10^-3^ ∼ 10^-7^ µg/ml) at 37 ℃ for 6h before infection. Intracellular viral RNA at 24 h post infection was quantified using qPCR. (**B**) Three other *Rhinolophus ferrumequinum* cell lines, including lung-origin RfPT, brain-origin RfBT and heart-origin RfHT were incubated with 10^-3^ µg/ml of indicated IFN proteins at 37 ℃ for 6 h, and then infected with HSV1 or PR8 viruses at an MOI=0.5 for 24 h. Intracellular viral RNA was detected using qPCR. IFN-treated or untreated RfKT (**C**) or RfBT (**D**) were fixed at 24 h post HSV1-GFP and PR8 (MOI=1) infection, viral positive cells were visualized by immunofluorescence assay. Data were presented as mean ± SEM. Error bar represents the standard error of three replicate experiments. The statistics were performed using one-way ANOVA with Dunnett’s T3 test. (*P < 0.05; **P < 0.01; ***P < 0.001; ****P < 0.0001; ns no significance, relative to the untreated).

The antiviral activity is largely determined by the extent or spectrum of ISGs induced by IFNs. Thus, we next compared the ISGs induced by the three IFNs. RfKT cells were treated with IFN proteins at the same dose, and ISG genes were determined using RNA-seq analysis. The data revealed that IFNaω induced a prominent higher number of ISGs (n=255) than IFNω (n=150) or IFNa (n=126) (Figure 6A and 6B). Moreover, IFNaω treatment resulted in not only a higher level of the common ISGs, but also a higher level of subtype-specific ISGs that may play important antiviral functions than the other two IFNs (Figure 6C and Table S4). This conclusion can be verified by qPCR targeting at three representative ISGs, *Mx1*, *OAS2* and *RASD2* (Figure 6D). Overall, IFNaω, the IFN that has the highest basal expression, also demonstrated the best antiviral activity or ISGs induction activity in *Rhinolophus ferrumequinum* bat.

**Figure 6.**
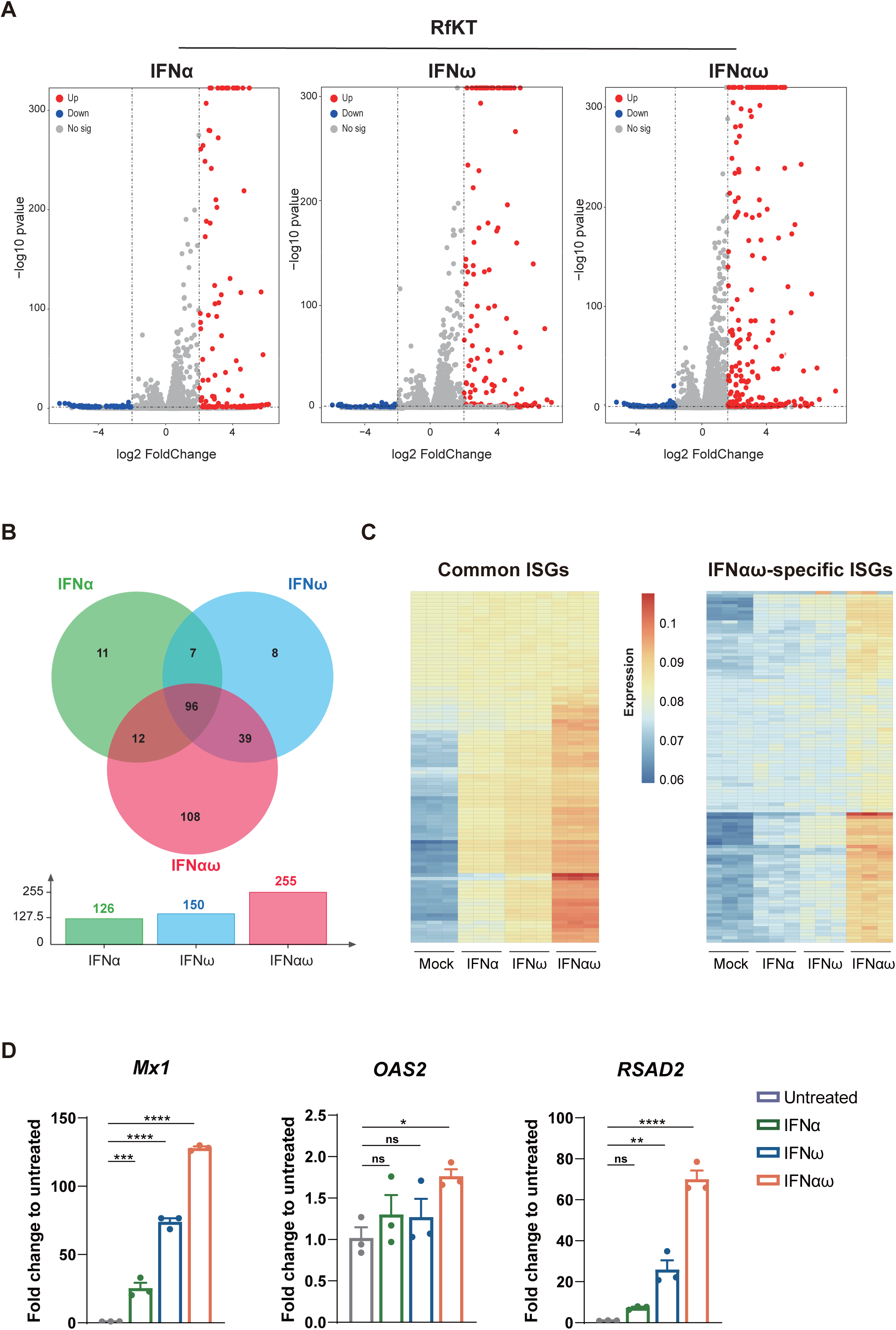
Differential induction of IFN-stimulated genes (ISGs) by *Rhinolophus ferrumequinum* IFN proteins in RfKT cell. (**A**) RNA-seq analysis of the RfKT cells treated with the indicated *Rf* bat IFN proteins (10^-3^ µg/ml) for 6 h. These data were subjected for gene differential analysis and shown as volcano plots. (**B**) Venn diagram comparing the ISGs induced by different IFN subtypes. (**C**) Normalized expression heatmap of the common or IFNaω-specific ISGs. (D) Induction of *Mx1*, *OAS2*, and *RSAD2* genes by different IFN subtypes in RfKT cells. Same dose of IFN were used (10^-3^ µg/ml for 6 h). Bars represent as mean ± SEM of n = 3 biological replicates. The statistics were performed using one-way ANOVA with Dunnett’s multiple comparisons test (**P < 0.01; ***P < 0.001; ****P < 0.0001; ns no significance).

### IFNαw protein demonstrated a higher thermo-stability and antiproliferative activity in Rhinolophus bat

Given that the body temperature of bats can reach up to 40 °C during flight, the thermo-stability of constitutively expressed IFNs is crucial to maintain a constitutive antiviral statement. Bat cells tolerate cell culture at a 40 °C without losing viability (18). In our experiment, different IFN proteins were incubated at 4 °C or 42 °C for 4 h, followed by a test for antiviral activity. Our data showed that a higher temperature affected the inhibitory ability of IFNs against viruses (HSV1 and PR8). Notably, the antiviral activity fluctuations of IFNω and IFNaω were minimal compared to IFNa (Figure 7A).

**Figure 7.**
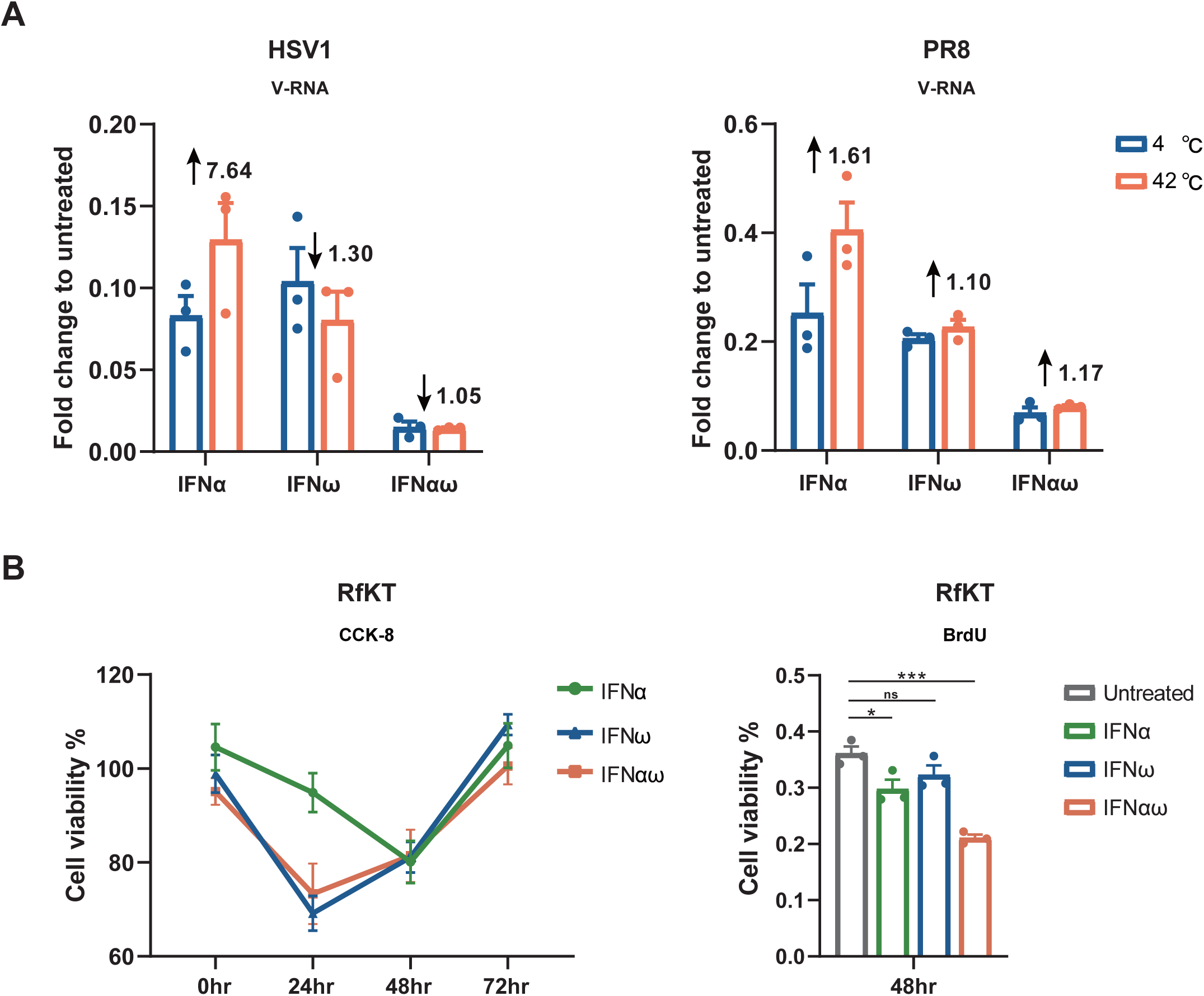
The thermo-stability and anti-proliferative activity analyses for bat IFN-I subtypes. (**A**) The thermo-stability comparison. The specified IFN proteins (10^-3^ µg/ml for each) were incubated at 4 ℃ or 40 ℃ for 4 hours, followed by incubation with RfKT cells at 37℃ for 6 hours. After a 24 hour infection, the expression levels of viral genes were analyzed using qPCR. The decreasing or increasing of V-RNA was shown as arrows. (**B**) Detection of the anti-proliferative activity of IFNs. Cell proliferation was measured by Cell Counting Kit-8 (CCK8) or BrdU assay measuring newly synthesized DNA in cells (0, 24, 48 or 72 h for CCK8, 48 h for BrdU). All experiments were repeated three times with similar results and a representative result is shown. One-way ANOVA with Dunnett’s correction was used for statistics (*P < 0.05; ***P < 0.001; ns no significance).

Finally, another important function for IFN-I protein is antiproliferation, which is often linked with apoptosis, tissue damage and anti-cancer activity (19). We then hypothesized that the constitutively expressed IFNs should also have constitutive antiproliferative activity. To test this, we drawn time-dependent antiproliferative curves of bat RfKT cells treated with IFNaω measuring live cells using CCK-8. The data indicated a shape decrease in cell viability upon IFNaω or IFNω treatments as early as 24 h, which is more severe than with IFNa treatment. To verify this result, we also used BrdU assay by measuring newly synthesized DNA. At 48h post treatment, IFNaω protein caused a significant reduction in cellular new DNA, whereas IFNa or IFNω proteins only caused mild changes (Figure 7B). Overall, the basal expressed IFNaω proteins not only demonstrated the best antiviral activity, but also showed the best antiproliferative activity or thermo-stability in *Rhinolophus* bat.

## Discussion

We hypothesized that the type I IFN system in bats may follow a different evolutionary path compared with other mammals, and this difference may contribute to a special coexistence between bats and viruses. Here, through a comprehensive analysis of the IFN genome locus or functional tests of IFN protein, we demonstrated a pan-bat dominance of *IFNω*-like responses in IFN-I system. In contrast to *IFNa*-dominated IFN-I loci in the majority of other mammals, bats generally have shorter IFN-I loci with more unconventional *IFNω*-like genes (*IFNω* or related *IFNaω*), but less or even no *IFNa* genes. In addition, bats generally have constitutively expressed IFNs, with the highest expressed more likely to be *IFNω*-like gene. In the case of *Rhinolophus* bat species, we confirmed that this highly expressed IFNω-like protein also demonstrated the best antiviral activity, anti-proliferative activity, and thermo-stability. Overall, bats choose a unique *IFNω*-like dominated IFN-I system, which is rarely found in other mammals. This difference provides insight into our understanding of how bats coexist with viruses.

We revealed general characteristics on bat IFN-I systems. There have been arguments around bat IFN-I system: (1) variability in the contraction or expansion of bat IFN-I systems depending on the species (3, 7); (2) most of the bat IFN-I studies are biased on knowledge from human IFNs that focused on *IFNa* genes (3, 7, 16). Our data indicated that there is a pan-bat contraction pattern in the IFN-I loci, as evidenced from shorter locus, lower number of genes and pseudogenization of *IFNβ* or *IFNε* genes in certain bat. A typical example is the entire lost of section between *KLHL9* and *IFNβ* for all Yangochiroptera bats. Similarly, our data also suggested that bats evolve to choose unconventional *IFNω*-like responses, which is different to most of other mammals (except armadillo). The gene expansion of *IFNω* is a signature event in bats and ungulates (15), but the dominance of *IFNω*-like genes in genome can only be found in bats and armadillos. An extreme example is the entire lost of *IFNa* genes in *Myotis* and *Pipistrellus* bats. Armadillo sits in the early stage of placental mammals’ evolutionary tree. Thus, it is conceivable that most bats maintained a rather ancient IFN-I system without extensive IFN gene expansion. Moreover, the constitutive expression of at least one IFN-Is has also been described in *Pteropus*, *Myotis* and *Rhinolophus* bats, but not in *Rousettus* bats (3, 7, 17, 20). Here, we showed that all six bat species maintained basal expression of IFN-Is, including the *Rousettus* bat. Thus, we conclude bat IFN-Is selected a very different evolutionary path to maintain a shorter locus, more ancient IFNs with constitutive antiviral or anti-proliferative activities.

This constitutive expression of *IFNω*-like proteins in bats appears to confer several advantages. limited analysis on human *IFNω* has indicated that the binding kinetics of *IFNω* to IFN-I receptors IFNaR1/R2 is faster than IFNa, suggesting which is more potent and faster ISGs inducer (21). This pattern is similar to *IFNβ*, which is important at quickly initiating downstream responses, but may not be fine-tuned as *IFNa* subtypes (22). Thus, it is conceivable that the IFNω-like proteins stimulate a constitutive antiviral statement in bats, this not only aids in early pathogen clearance but also primes the down-stream IFN responses by other IFN-I subtypes if the viruses proliferate to high titer. Furthermore, this thermo-stable IFNω-like protein ensures the maintenance of antiviral activity under different physiological conditions, particularly relevant given the elevated body temperatures bats experience during flight. Lastly, bats are known to have a relative long lifespan (23). As cancer is one of the major threats of human longevity, the constitutive anti-proliferative activity of IFNω-like proteins in bats may play a role in suppressing tumor growth. Overall, we believe the unique bat IFN-I system is related to its longevity or as reservoir hosts of viruses, albeit the mechanisms behind this link are not fully understood.

There are still many unknown about bat IFN-I systems, with one major question is the cellular driver of these constitutively expressed IFNs. A list of known human IFN-regulatory factors, including *IRF1*, *IRF3* or *IRF7* has been tested in bats. Apparently, the regulator of basal IFNs in bats are an atypical factor yet to be identified, but not these known IRFs (24–26). Likewise, what is the physiological driver of a basal IFN expression is another question to be answered. As the only flying mammals, bats were believed to possess an increased metabolic capacity (6). Whether a higher metabolic rate or the byproducts of oxidative metabolism (such as reactive oxygen species, ROS) induced basal IFN expression needs to be discovered in future studies (6). Finally, we realized that pangolin is the only evolutionarily close mammal that shares a similar short IFN-I locus with bats. Coincidently, pangolins are also believed as important intermediate hosts for bat SARS-or MERS-related coronaviruses spillover. Different to bat IFN-Is, pangolins have more *IFNa* and a pseudotyped *IFNε* gene, which may dampen the innate immunity against viruses (27). Whether an *IFNω*-dominated response in bats is superior to an *IFNa*-dominated response in pangolin is another questions to be answered.

In summary, we demonstrated pan-bat characteristics in type I IFN system, including a dominance of *IFNω*-like genes in genome locus and a dominance of constitutive antiviral IFNω-like responses. The study provides insight into our understanding of the innate immunity that contribute to a special coexistence between bats and viruses.

## Acknowledgements

We thank Prof. Yan-Yi Wang, Min-Hua Luo and Quan-Jiao Chen from Wuhan Institute of Virology, Chinese Academy of Sciences for their support of viral resources. We thank Prof. Lin-Fa Wang from Singapore Duke-NUS Medical School for the support of the SV40LT plasmid.

## Data availability

All data relevant to this study are included in this article (and its supplementary information file).

## Footnotes

The work was supported by the China National Science Foundation for outstanding scholars (82325032 to P.Z.) and the Strategic Priority Research Program of the CAS (XDB29010204 to P.Z).

