## Supplementary Figures and Tables for "Unconventional *IFNω*-like genes dominate the type I IFN locus and the constitutive antiviral responses in bats": 2-Supplementary Figures.docx

**Supporting Information**

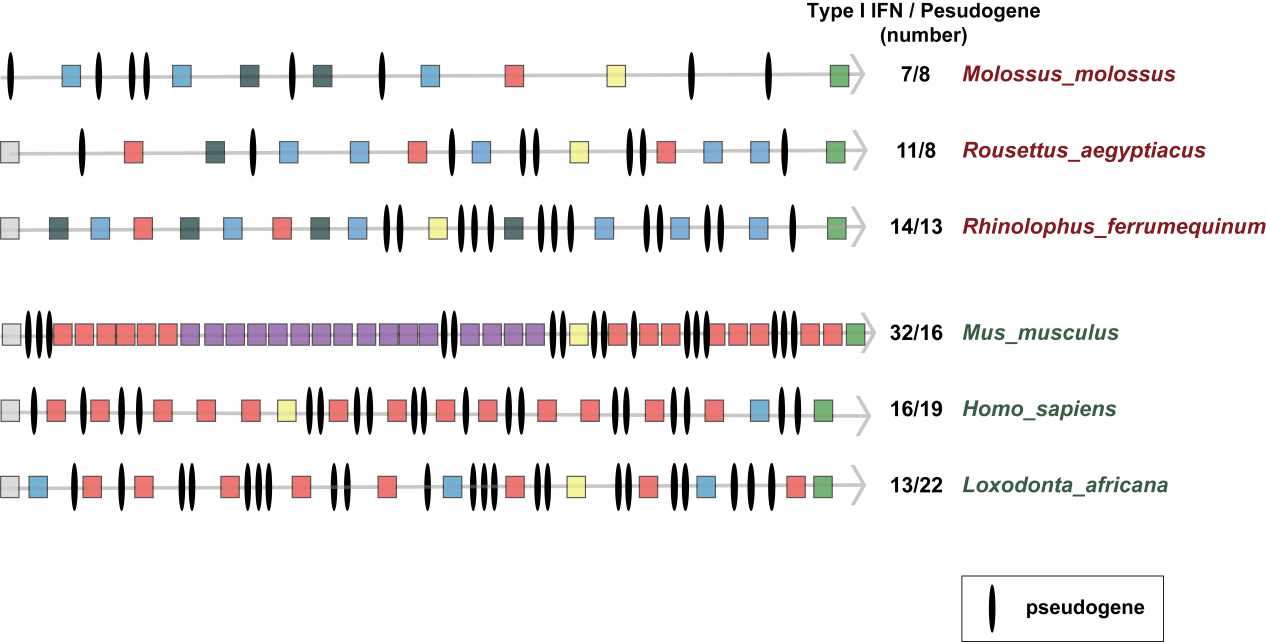

**Figure S1 Characterization of pseudogene in IFN-I locus.**

Three bats and three non-bat mammals were subjected for the analysis. Black lines represent pseudogenes. Intact ORFs were not shown.

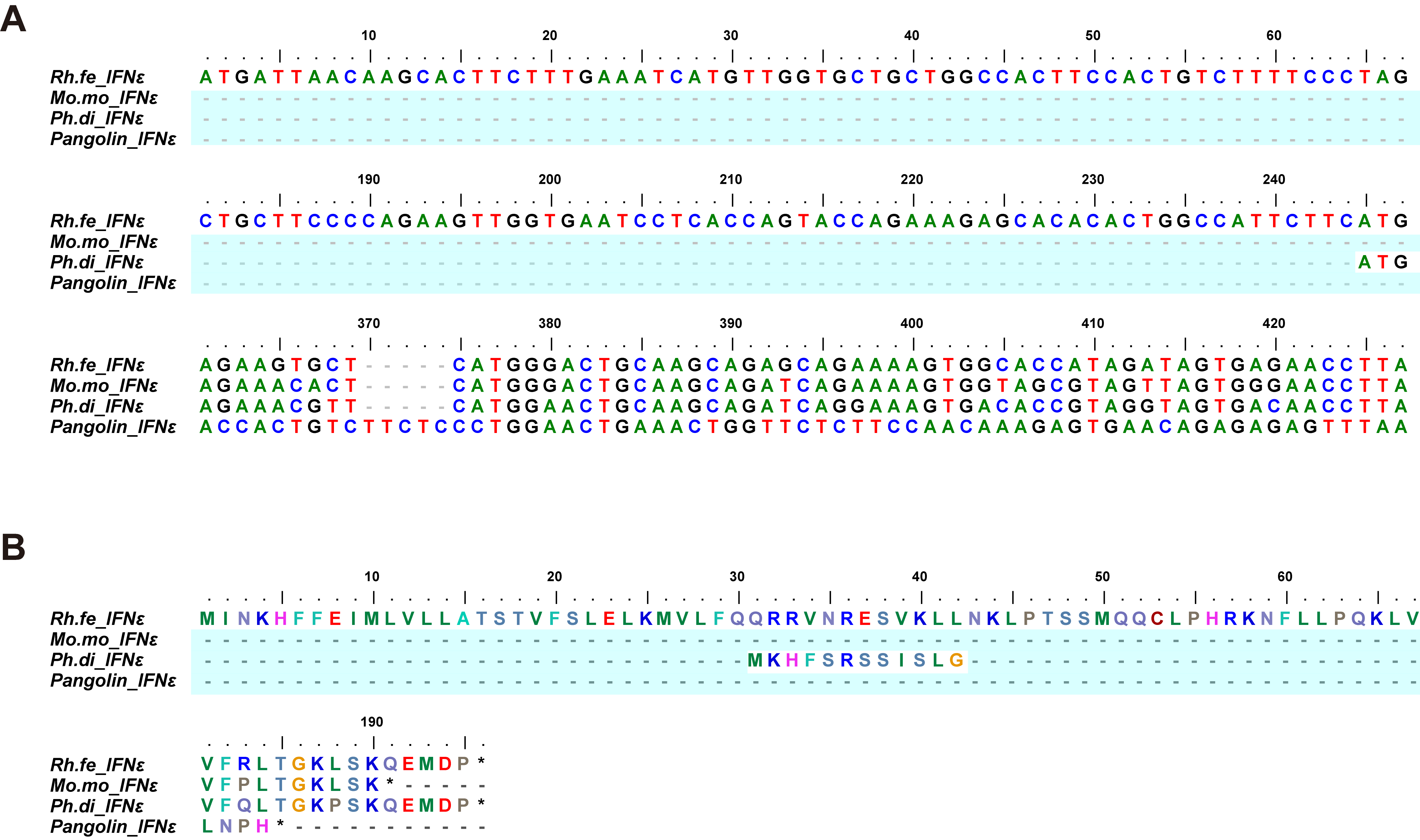

**Figure S2** **Sequence comparison of *IFNε* pseudogenes*.***

(**A**) Nucleotide alignment of *IFNε* genes from *Rhinolophus ferrumequinum, Molossus molossus, Phyllostomus discolor* and pangolin. (B) Amino acid sequence alignment of *IFNε.* Dashed lines indicate deletions. Compared to *Rhinolophus ferrumequinum*, other three *IFNε* genes contain large deletions.

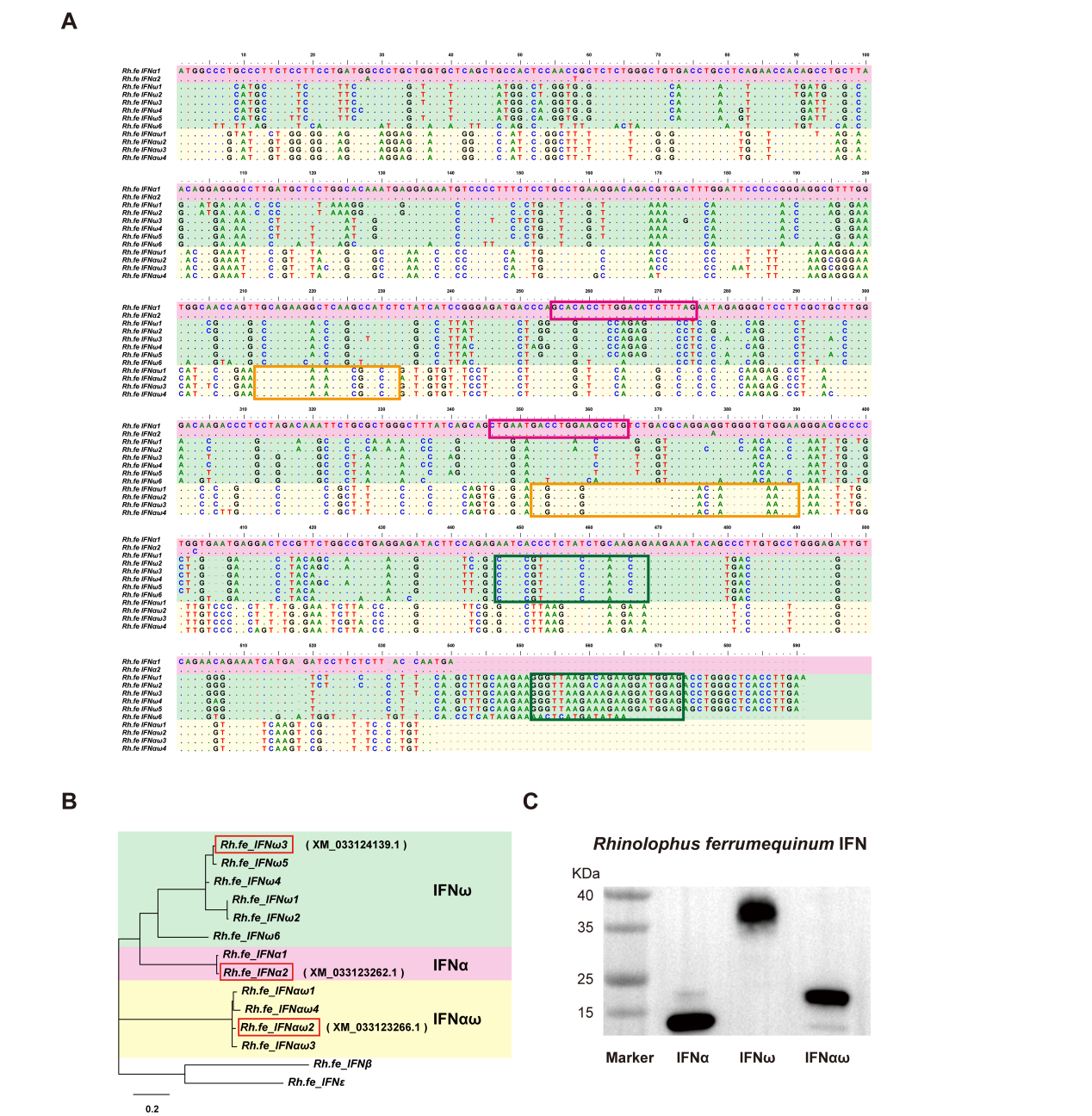

**Figure S3** **Sequence analysis of IFN-Is in *Rhinolophus ferrumequinum.***

(A) Alignment of *Rhinolophus ferrumequinum* IFN-Is. The pan-specific of primers used for different IFN subtypes was labeled with boxex. (B) Phylogenetic analysis for *Rhinolophus ferrumequinum* IFN-Is*. Rhinolophus ferrumequinum* type I IFN used to express proteins in this study are highlighted in red boxes, the sequence numbers of these three IFN-Is are in bracket. (C) Rf-IFN proteins were overexpressed using a mammalian cell expression system (HEK293F). Concentrated IFNs were subjected to western blotting to verify protein purity. Note that the bands of IFNω have apparent molecular weights higher than expected values (~ 20 kDa), which might reflect posttranslational modification such as N-glycosylation (28).

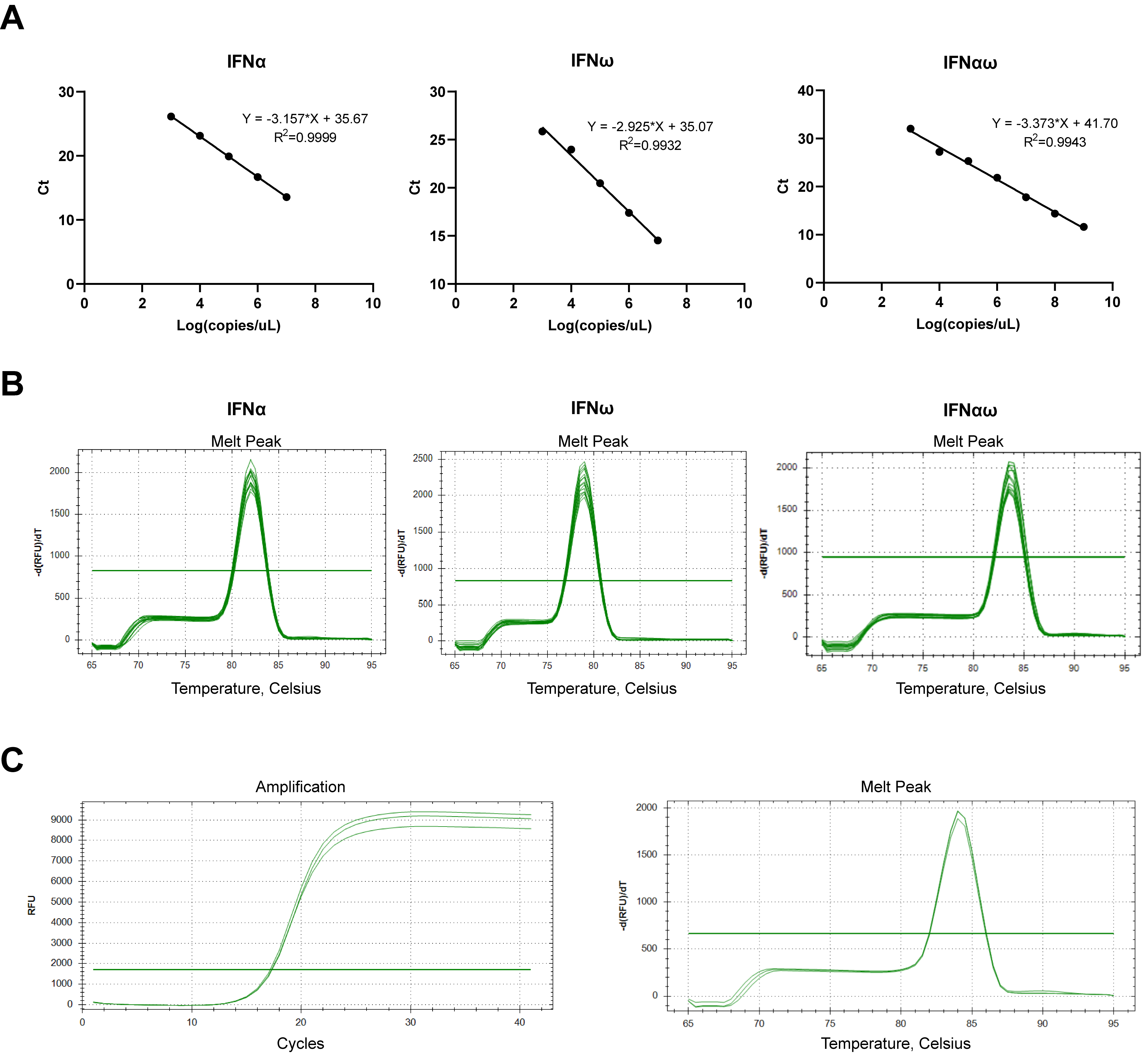

**Figure S4 Efficiency and specificity of *Rf* qPCR specific primers**

(A) The standard curve (B) the melt peak of the three Rf-IFN subtypes qPCR primers. (C) Amplification curve and melt peak of the reference gene *β-actin* in *Rhinolophus ferrumequinu*m. The graph on the left showed the amplification curve of this gene in *Rf* bat cells under different stimulations (Mock vs IFN-treated vs Infection), with a Ct value stable at around 17 copies. The image on the right showed that the Tm value of this gene was 84 °C.
